# Carbon acquisition in a Baltic pico-phytoplankton species - Where does the carbon for growth come from?

**DOI:** 10.1101/2020.09.07.285478

**Authors:** Luisa Listmann, Franziska Kerl, Nele Martens, C-Elisa Schaum

**Affiliations:** University of Hamburg, Institute for Marine Ecosystem and Fisheries Science, 22767 Hamburg; Centre for Earth System Science and Sustainability, 20146 Hamburg

**Keywords:** Pico-phytoplankton, environmental change, carbon acquisition, evolutionary history, primary production

## Abstract

- Pico-phytoplankton have ample scope to react to environmental change. But we know little about the underlying physiological mechanisms that govern how evolutionary history may affect short-term responses to environmental change.
- We investigated growth rates and carbon uptake related traits (i.e. fitness proxies) in different temperatures and at different times during the microbial growth curve of eight novel strains of *Ostreococcus sp*. (ca. 1-2µm). The strains were isolated from two distinct regions of the Baltic Sea differing in salinity and temperature from North-East (Bornholm Basin) to South-West (Kiel area).
- Strains from the warmer, more variable Kiel area had higher growth rates in general and showed more variable growth rates compared to strains from the colder and less variable Bornholm Basin.
- In addition, growth was maintained in early stages of the growth curve by organic carbon acquisition and the increase in growth over time and with temperature was associated with an increase in inorganic carbon acquisition (net primary productivity).
- Based on the differences between net primary productivity and potential growth on organic carbon, we postulate a shift in carbon acquisition between inorganic and organic sources in *Ostreococcus* sp. with potential implications on ecological dynamics within microbial communities.

## Introduction

In recent years, temperatures in the atmosphere and sea surface have been increasing at an unprecedented rate, putting organisms into environmental conditions that they have likely not experienced before (*IPCC Fifth Assessment Report - Synthesis Report*, 2014). An organism can react to a changing environment through a combination of strategies such as moving to other locations, coping with changes *via* plasticity (i.e. phenotypic variation within one genotype), adapting (i.e. evolutionary change that leads to an increase in fitness) or in the worst case, die (Gienapp *et al*., 2008). Here, we investigate the adaptive response (i.e. how growth rates differ with respect to temperature in the short-term within several generations) of a pico-phytoplankton species *Ostreococcus* sp. from the Baltic Sea. Growth rates as fitness proxies can indicate the direction and magnitude of change in fitness (Elena & Lenski, 2003). In addition to understanding if short-term responses to warming in *Ostreococcus* vary, we are also interested in *how*. Thus, we also want to mechanistically link evolved growth responses to the underlying metabolic responses.

Pico-phytoplankton are a globally distributed group of microbial photosynthetic primary producers that make up about 0.53–1.32 Pg C of the marine biomass and contribute to ca. 20% of marine primary production (Worden *et al*., 2004; Buitenhuis *et al*., 2012). In coastal areas, pico-phytoplankton can temporarily contribute up to 80% to the marine production of oxygen via photosynthesis and fuel biogeochemical cycles (Worden, 2006). As primary producers, they assimilate inorganic carbon (CO_2_) and have important cascading effects on higher trophic levels in the marine food web (Field *et al*., 1998; Falkowski *et al*., 2008). In addition, high growth rates and large populations size with high standing genetic variation (Reusch & Boyd, 2013) enable pico-phytoplankton, as well as other microbial species, to track today’s rapid changes in the environment *via* fast plastic responses and evolutionary change and often through a combination of both (Lenski & Travisano, 1994; Wiser *et al*., 2013; Levis *et al*., 2016).

Previous experimental long-term studies have shown that phytoplankton are indeed able to adapt to environmental change within a few hundred generations which translate to several months to years in the laboratory (Lohbeck *et al*., 2012; Schlüter *et al*., 2014; Listmann *et al*., 2016; Schaum *et al*., 2018). Despite their insights into the adaptive potential of phototrophic microbes, these experiments focus mainly on single strains (i.e. genotypes) of species from culture collections and are very time consuming and large experiments. As a result, they only capture a drastically reduced image of ecological variability. The latter, however, may add to the characterization of the adaptive potential of organisms, because with ecological variation, the range of phenotypes within a species, that essentially selection can act on, increases (Des Roches *et al*., 2018). In this study we circumvent the limitations of long-term experimental studies on laboratory strains by using two approaches: First, to account for a degree of ecological variability (Boyd *et al*., 2013; Hattich *et al*., 2017; Godhe & Rynearson, 2017), we use not one, but eight strains of the same species complex. Second, in order to investigate the evolved response to warming, we use strains of the same species complex with different environmental histories - an approach called space for time substitution (Likens, 1989).

To understand *how* pico-phytoplankton evolve to warming, we want to link growth responses to the underlying metabolic responses (e.g. Padfield *et al*., 2016). Metabolic responses that are associated with growth can include nutrient uptake related strategies (Sommer, 1984; Edwards *et al*., 2015), metabolic responses within the cell for energy turnover or allocation (Rokitta *et al*., 2016; Collins & Schaum, 2019) or, as is most important for primary producers, carbon uptake related strategies (Rost *et al*., 2006). Several studies on different phytoplankton groups have demonstrated that net primary production, which describes the uptake of carbon for growth, can change in response to changes in the environment essentially modulating the relationship of net primary production and growth (Schaum *et al*., 2017b; Barton *et al*., 2020). These indicate that inorganic carbon is assimilated in different quantities. However, these studies have so far mainly investigated the responses in cultures at exponential phase in the microbial growth curve. This allows us to understand metabolic dynamics that are associated with exponential growth, but it ignores that ample theory in ecology and evolution would predict the existence of multiple strategies both depending on the environmental condition and the life cycle state of a microbial organism (Halsey *et al*., 2013; García-Carreras *et al*., 2018). Here, we additionally focus on metabolic responses during early and late exponential growth phases rather than only the maximum exponential growth phase of the microbial growth curve.

Phytoplankton species can be divided into different functional groups regarding their carbon uptake. Of these, photoautotrophs assimilate carbon via photosynthesis while mixotrophic phytoplankton acquire their carbon either via photosynthesis or uptake of organic carbon compounds (Rebecca Lindsey & Scott Design by Robert Simmon, 2010). In this study, we focus on a globally distributed pico-phytoplankton species, *Ostreococcus* sp., which has a size of about 1µm and is the smallest known free-living eukaryote (Courties *et al*., 1994; Rodríguez *et al*., 2005). *Ostreococcus* sp. is characterized as a photoautotrophic species, i.e. using CO_2_ as its carbon source for growth (Courties *et al*., 1994). However, it has been shown that *Ostreococcus* has the potential to grow in the dark relying on other carbon sources (e.g. sorbitol) that do not directly come from co-occurring photosynthesis (van Ooijen & Millar, 2012). Therefore, in *Ostreococcus* other carbon uptake related strategies could play a role to sustain growth. Here, we want to specifically test whether and under which conditions uptake of organic carbon occurs in the light. In other words, instead of solely investigating how much carbon is allocated to growth, we study where the carbon for growth is coming from, considering effects of changes in the thermal environment (temperature assay), evolutionary history (via space for time substitution), and the life cycle (throughout a growth curve). In addition, we investigate how much intraspecific variation exists in carbon uptake related strategies.

We successfully isolated eight novel strains of *Ostreococcus tauri* (7) and *Ostreococcus mediterraneus* (1) in Spring of 2018 (RV ALKOR cruise AL505) (Table 1) from the Baltic Sea in order to study a range of strains of the same species complex with different evolutionary histories. The Baltic Sea is characterized by different environmental gradients including for example temperature, salinity or nutrients (Leppäranta, Matti, 2009) that have changed in the last decades (Zhong *et al*., 2020). Here, we focus on two regions in the South-West and East of the Baltic, the Kiel area and Bornholm Basin, differing mainly in temperature and salinity. The respective gradients range from warmer, more variable temperatures and higher salinity in the Kiel area compared to colder, less variable temperatures and lower salinity in the Bornholm Basin. On the new strains of *Ostreococcus* sp., we measured the growth response to two different temperatures and how the respective evolutionary trajectories affected this response. In addition, by quantifying net primary production *via* photosynthesis and respiration measurements (Schaum *et al*., 2017a) and potential growth on organic carbon substrates (Hackett & Griffiths, 1997; Rutgers *et al*., 2016), we characterized different strategies of how growth can be maximized (or changed) in varying environments and at different time-points during the exponential growth phase.

**Table 1:**
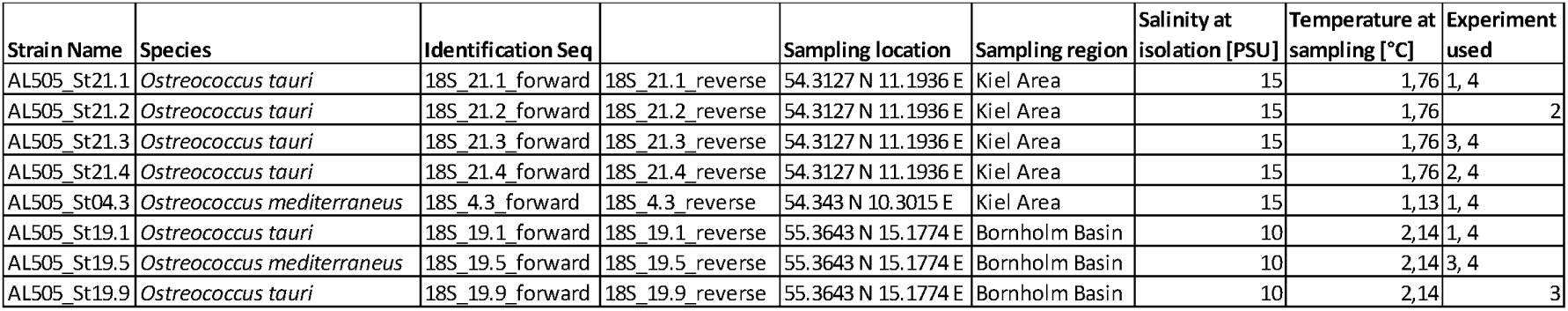
This table summarizes the different origins of the strains that were used in the experiments. The given parameters of isolation were taken on board the research vessel at the time of sampling of the phytoplankton community from which the *Ostreococcus* strains were isolated. The sequences for identification via 18SrRNA are uploaded as supplementary data.

## Material and Methods

### Ostreococcus isolation and experimental set-up

We isolated *Ostreococcus* sp. from pico-phytoplankton community samples obtained during a RV ALKOR cruise (AL505) in 2018 (see Fig. **1a** and Table 1 for sampling dates and locations) using a Niskin bottle at 5m. Community samples were immediately passed through a 35µm sieve to remove grazers and large debris, and then further size fractioned *via* gentle filtration through a 2µm membrane filter (kept filtrate) and a 0.2µm filter (kept filter and rinsed gently). From these samples, we successfully isolated eight new strains of *Ostreococcus* sp. (see Table 1 for details); five from the Kiel area and three from the Bornholm Basin.

**Figure 1.**
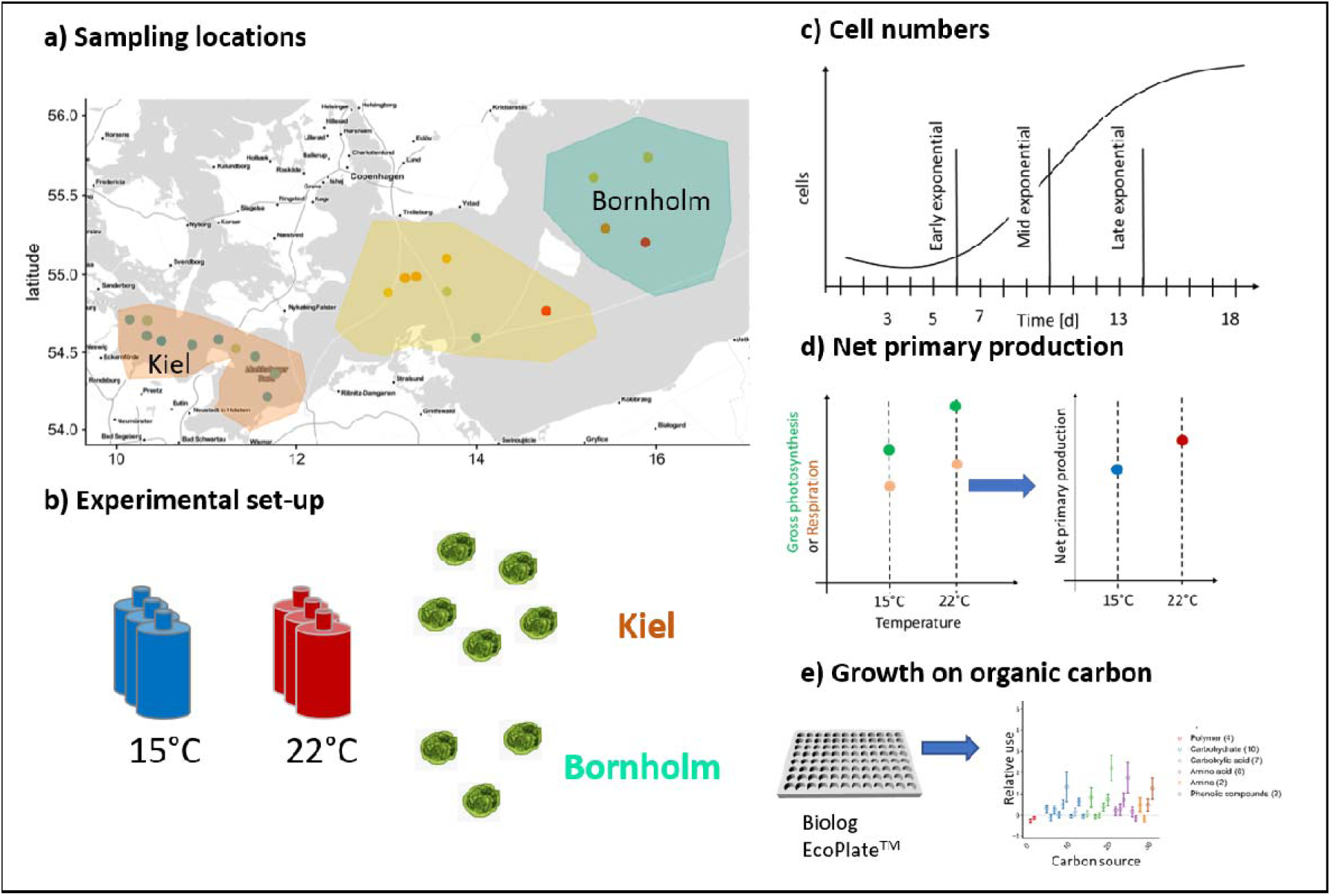
Sampling locations and experimental set-up: The pico-phytoplankton communities from which we isolated *Ostreococcus* sp. were collected during two research cruises in March and August 2018 and originate from Kiel and Bornholm area (panel **a**). Eight successfully isolated strains of *Ostreococcus* sp. were exposed to 15°C and 22°C (panel **b**) in four consequent experiments and monitored daily for growth *via* tracking cell numbers (panel **c**). In addition, we measured net primary production daily (panel **d**) in experiment 1-3 and in experiment 4 we investigated potential growth on organic carbon at three time points of the microbial growth curve (panel **e**).

The eight new strains were cultured to determine their growth rate and metabolism response to 15°C and 22°C (spanning late spring and late summer temperatures). The experiment had to be carried out in three subsequent batches due to the limited number of metabolic measurements possible at the same time (see Fig. **1b**, Table 1). The batches were all set up the same way: each strain was replicated three times and each replicate inoculated with 3000 cells/mL in 40mL f/2 media (Guillard, 1975) of the respective salinity of isolation (Table 1). All replicated cultures were exposed to the two treatment temperatures with a 12:12 day and night cycle at 100µE light intensity for 18 days ensuring growth through a whole microbial growth cycle (Fig. **1c**). Starting at day three of microbial growth we measured cell numbers daily *via* flow cytometry (BD Accuri C6 flow cytometer) and starting on day four to five, we measured photosynthetic metabolic activity daily *via* optical O_2_ measurements (Fig. **1d**).

Due to logistic limitations, for the measurement of growth on organic carbon sources we set up an additional experiment using six representative strains of the eight isolated *Ostreococcus* strains (four from Kiel and two from Bornholm, respectively) (Fig. **1e**). Each strain was inoculated at 3000 cells/mL in 200 mL of f/2 media (Guillard, 1975) of the respective salinity of isolation and exposed to both 15°C and 22°C with a 12:12 day and night cycle at 100µE light intensity. At three time points during the microbial growth curve (determined via the preceding growth experiments), we investigated the potential of each strain to grow on 31 different organic carbon sources using ecoplates (Biolog EcoPlate™).

### Determination of growth rates and net primary production in experiments 1-4

On the daily cell count measurements, we fitted a growth curve containing a lag phase, exponential phase and carrying capacity. To analyse the shape of the growth curves, non-linear curve fitting of a gompertz growth model (Buchanan *et al*., 1997) was carried out using the ‘nlsLoop’ function in the R package, ‘nlsLOOP’ (version 1.2-1). Parameter estimation was achieved by running 1,000 different random combinations of starting parameters for cell count at carrying capacity, duration of lag phase, and maximum growth rate picked from a uniform distribution. The script then retained the parameter set that returned the lowest Akaike information criterion (AICc) score, yielding *µmax* and *day at µmax*. In addition, we calculated growth rates at early and late exponential phase (three days before and after day at µmax, respectively) using the following formula (ln(N_t_-N_t-1_)) with N being the number of cells/ml. The *day at µmax* was important for subsequent analysis of growth on organic carbon sources (see Table S1).

Net photosynthesis and respiration rates were measured on PreSens ® SDR Sensor Dish optodes. We measured oxygen production for 15 minutes in the light, and respiration for 15 minutes in the dark under the light-and temperature conditions set in the incubator (i.e. all experimental units at their assay temperatures). All characterizations were carried out at the same time of day (9am to 11am). From these measurements we calculated net primary production rates (following (Falkowski *et al*., 1985) in µgC*cell^-1^*day^-1^).

### Potential growth on carbon via ecoplates

Here we were interested, if and how much a culture could grow on a number of different carbon sources that we provided via the ecoplates. Based on Exp. 1-3 we identified three time points (corresponding to three different days during the microbial growth cycle) at which we then measured growth on organic carbon. These time points were “early exponential phase” three days prior to day at µmax, “mid exponential phase” at µmax and “late exponential phase” three days later than day at µmax. The actual days of inoculation into ecoplates varied between the different strains of *Ostreococcus* and treatment temperatures (see Table S2 for details). Each ecoplate contained 31 different organic carbon sources in triplicates and three controls with water (see Rutgers *et al*., 2016 for list of sources and groups thereof and Fig. S1). Each of the 96 wells was inoculated with 200µl of culture (except one control of water with only MQ) and then left to grow for 24h at the respective experimental condition as the original culture. After 24h we fixed the samples with sorbitol (3µl in 200µl sample) for 24h at 4°C in the dark and then froze them for later analysis via flow cytometry. After thawing the ecoplates overnight we counted the cells in each well and calculated the relative change in cell numbers on organic carbon compared to the control (on water). In addition, we calculated an overall value of relative increase in cell numbers on organic carbon compared to a control without the addition of organic carbon the following way: First, we calculated the mean of cells/mL for the control wells that only contained water; second, we calculated the relative change in cells/mL for each of the 93 wells that contained a carbon source compared to the control (i.e. “water”, no other organic carbon source). Third, we counted the number of wells where the relative change in cell numbers was positive. And last, the overall potential growth on organic carbon was then calculated as the mean of the relative increase of cells normalized by the number of wells on which the relative increase was positive.

### Statistical analysis

All data were analysed in the R programming environment (version 3.6.3.) using the following packages ‘nlme’, ‘ggpot2’, ‘lme4’, ‘emmeans’, ‘vegan’, ‘reshape2’, and ‘multcomp’.

All experimental responses (µmax, growth rate at early and late exponential phase, net primary production and overall growth on organic carbon) were analysed with a linear mixed effects model (lme) (within the nlme package, version 3.1-137). We focused on the effect of timing (i.e. differences between early, mid and late exponential phase in the microbial growth curve) at 22°C and the effect of temperature in the mid exponential phase since we had all experimental responses for these time points and temperatures. The responses were first analysed *via* a *global model* (Table S5-7 supplements) that included sampling location (Kiel area or Bornholm Basin) and assay temperature (15°C and 22°C) or time (for the responses at early, mid and late exponential phase) as fixed factors in full interaction. The “experiment” (Exp 1-3 for µmax and net primary production) was computed as a nested random effect within region. We subsequently analysed the growth rate again for each region separately to further characterize the intra-regional variation of responses at mid exponential phase and in response to temperature (*regional model* Table S5supplements). The regional model included strains (five different ones in the *Kiel regional model* and three different ones in the *Bornholm regional model*, respectively) and assay temperature (15°C and 22°C) as fixed factors in full interaction. All lme models were reduced to the single and interacting factors containing the lowest AICc score with a minimum difference of 2.

The changes in cell numbers on the organic carbon compounds was analysed in two ways: First we analysed the differences in the groups of carbon compounds via a linear mixed effects model containing the effect of timing at 22°C (see above) and temperature in mid exponential phase as well as the effects of region and “carbon group”. The lme model was reduced to the single and interacting factors containing the lowest AICc score with a minimum difference of 2. Second, we confirmed the statistical analysis on the relative growth on carbon and changes in cell numbers via a PCA analysis and subsequent permanova that tested again for the effect of timing at 22°C (see above) and temperature in mid exponential phase. The difference to the first analysis is, that it includes the differences between all 31 carbon sources and how the complete use of all the sources differed between time-points, temperature and regions.

## Results

### Adaptive response in Ostreococcus measured via growth rates

Following the expected shape of a microbial growth curve, the highest growth rates were reached in the middle of the microbial growth curve and lower (down to no growth) at the early stage of the microbial growth curve both in the Kiel and Bornholm strains (Fig. **2**, Table S5 *Global model* (TI) Effect of “Timing” F_2_=4.990, p=0.011). Towards the end of the exponential phase, growth rate decreased again but not significantly (contrast mid to late exponential phase t-ratio= 1.704, p=0. 5357). In the Kiel strains, the growth rates were higher in all three phases compared to the Bornholm region (Fig. **2**, Table S5 *Global model* (TI) Effect of “Region” F_1_ =11.779, p=0.001). All strains of *Ostreococcus* increased their growth rate from 15°C to 22°C (Fig **2**, Table S5 *Global model* (TE) Effect of “Temperature” F_1_ =40.162, p<0.0001), however, this increase varied strongly between the strains within the Kiel region (Table S5 *Regional model* (Kiel) Effect of “Temperature” F_1_ =31.725, p=0.0001; Effect of “Isolate” F4 =7.618, p=0.003) and between the two regions (Table S5 *Global model* (TE) Effect of “Region” F_1_ =15.430, p=0.0005). This indicated an effect of the absolute difference in experienced past environment including the variability therein; in other words, their evolutionary history.

**Figure 2.**
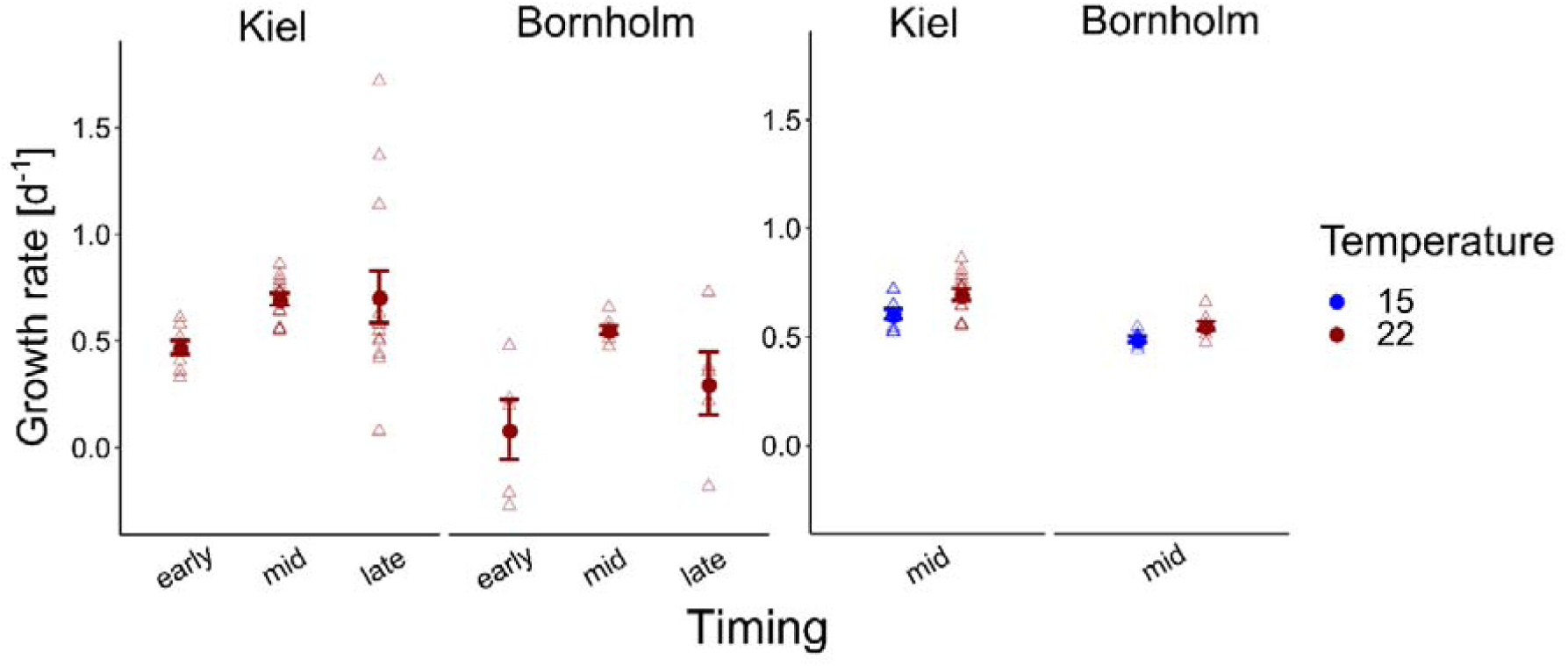
Growth rates are shown here at 22°C (red) at early, mid and late exponential phase in the left panel and at 15°C (blue) and 22°C (red) at mid exponential phase in the right panel. Note that for the growth rate at mid exponential phase we used a logarithmic curve fit to all numbers collected during the experiment whereas early and late exponential growth rates were estimated via ln(N_t_-N_t-1_) three days prior and after the day where growth was maximum. All points show mean +/-1 SE including n=15 and n=9 for Kiel and Bornholm, respectively. Triangles present the growth rate of each experimental unit.

### Inorganic carbon acquisition

The net primary production in both Kiel and Bornholm strains increased with temperature at mid exponential phase (Fig. **3**, Table S6 *Global model* (TE) Effect of “Temperature” F_1_ =8.818, p=0.0005) and from early to mid exponential phase at 22°C as well (Fig **3**, Table S6 *Global model* (TI) Effect of “Timing” F_1_ =36.583, p<.0001). From mid to late exponential phase there was no significant increase anymore (contrast mid to late exponential phase t-ratio=-0.398, p=0.916). At the mid exponential phase, net primary production was slightly but not significantly higher in the Bornholm strains compared to the Kiel strains both at 22°C and 15°C (Fig. **3**, Table S6 *Global model* (TE) Effect of “Region” F_1_ =1.999, p=0.261). At the early exponential phase at 22°C net primary production was, however, lower in Bornholm (Figure 3, Table S6 *Global model* (TI) Effect of “Timing * Region” F_1_ =5.093, p=0.006). Net primary production increased in most cases because GP became relatively higher compared to R (supplementary Fig. S3).

**Figure 3.**
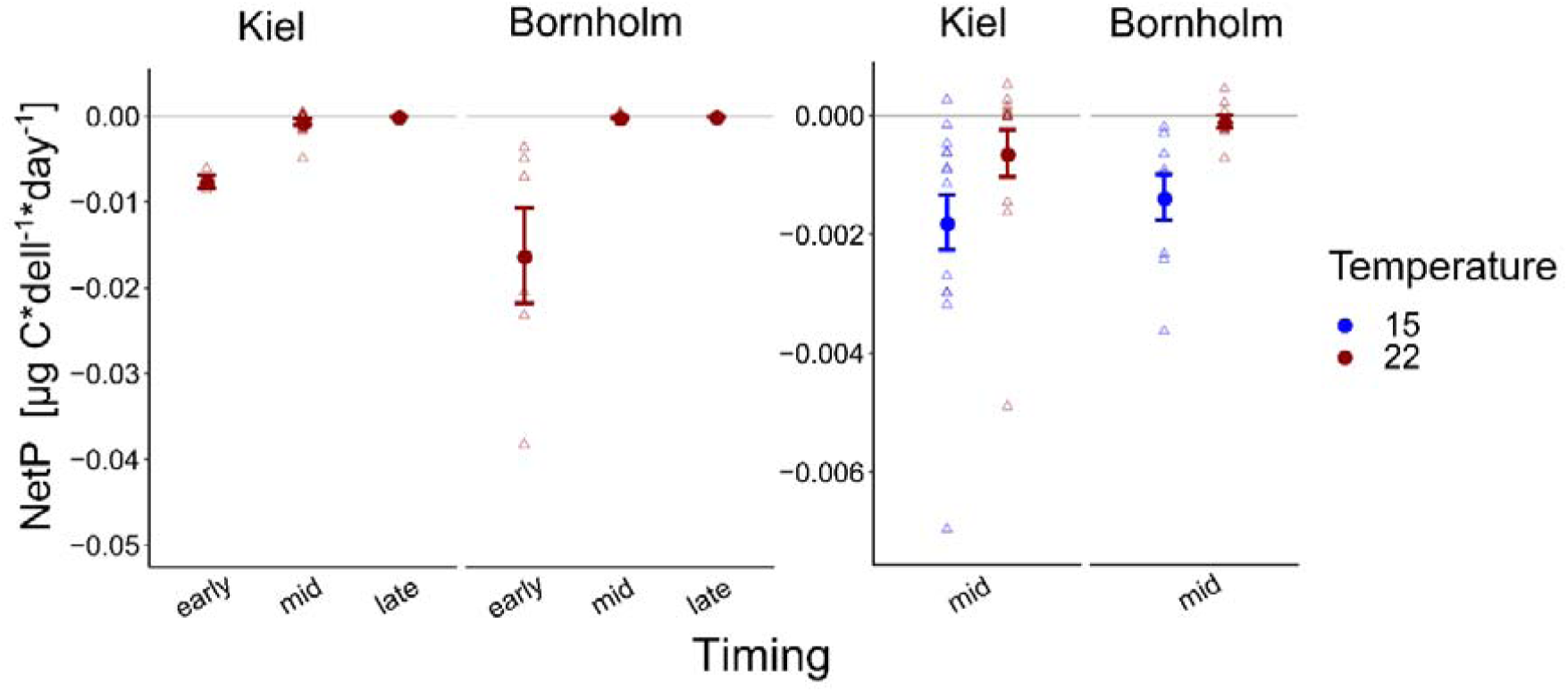
Net primary production (NetP= is shown here at 22°C (red) at early, mid and late exponential phase in the left panel and at 15°C (blue) and 22°C (red) at mid exponential phase in the right panel. Note that the axes are different between the panels due to the strong increase in net primary production from early to mid exponential phase. All points show mean +/-1 SE including n=15 and n=9 for Kiel and Bornholm, respectively. Triangles present the net primary production of each experimental unit.

### Growth on organic carbon sources and correlation to net primary production

Due to experimental limitations, we were only able to measure growth on organic carbon sources on one representative replicate of four strains from the Kiel area and two strains from the Bornholm area, respectively. Thus, a comparison between the strains was statistically not possible. Averaged over all the organic sources we did find a higher use of organic carbon sources by isolates from Kiel compared to Bornholm at mid exponential phase in both assay temperatures (Fig. **4**, Table S7 *Global model* (TE) Effect of “Region” F_1_ =11.416, p=0.027) as well as early and late exponential phase at 22°C (Fig. **4**, Table S7 *Global model* (TI) Effect of “Region” F_1_ =6.531, p=0.063). In addition to the overall difference between the regions we also found effects of the timing of measurement and temperature: on the one hand, the growth on organic carbon in both Kiel and Bornholm strains decreased from early to mid and late exponential phase at 22°C (Fig. **4** Table S7 *Global model* (TI) Effect of “Timing” F_2_ =12.398, p=0.002). On the other hand, at mid exponential phase, the growth on organic carbon sources was higher at 15°C compared to 22°C in both areas (Fig **4**, Table S7 *Global model* (TE) Effect of “Temperature” F_1_ =5.730, p=0.062).

**Figure 4.**
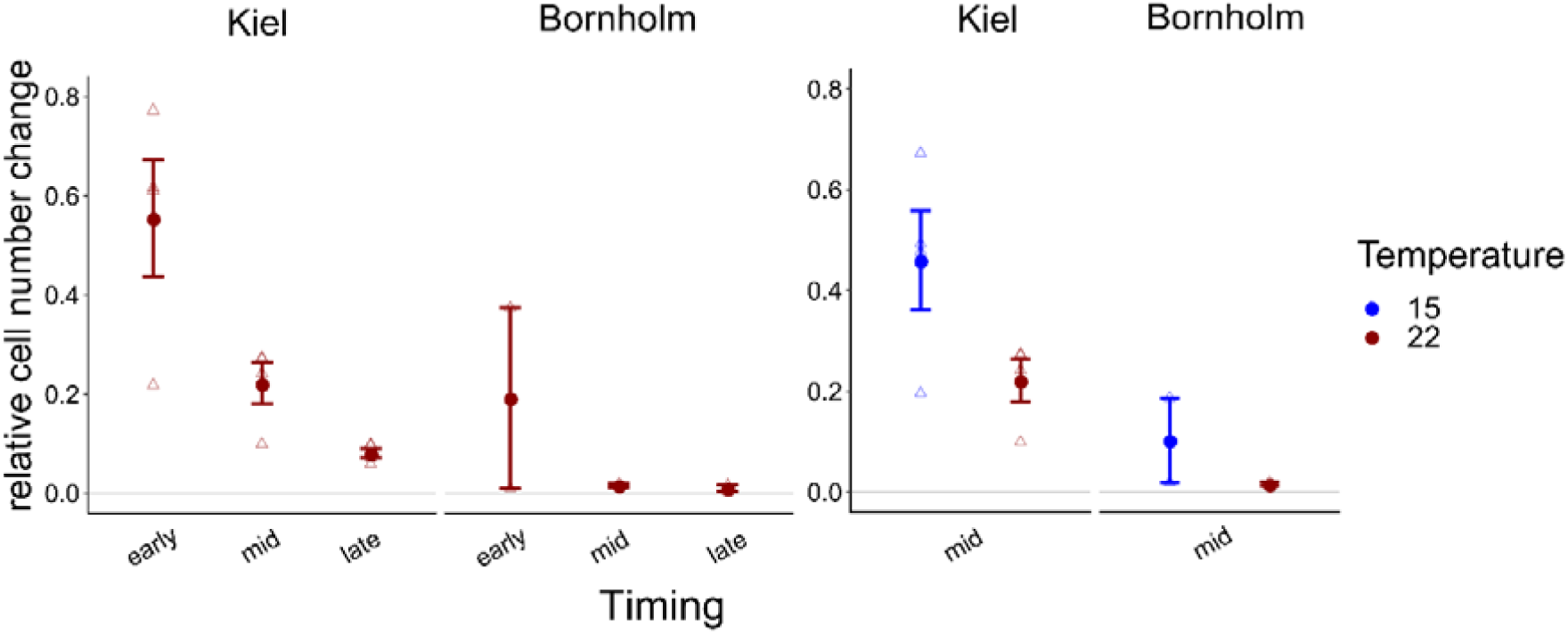
Overall relative increase in cell numbers on carbon normalized by the number of sources on which we found positive growth is shown here: in the left panel the growth at 22°C (red) at early, mid and late exponential phase is shown, whereas in the right panel relative growth at mid exponential phase at 15°C (blue) and 22°C (red) degrees is shown. All points show mean +/-1 SE including n=4 and n=2 for Kiel and Bornholm, respectively. Triangles present the overall growth on organic carbon per ecoplate of each experimental unit.

In addition to the differences in growth when all carbon sources were considered together, we found that at different sampling times and temperatures the carbon sources were used in different quantities (Fig. **5**). In the early exponential phase on all carbon groups except for the polymers (Cyclodextrin and Glycogen, see Fig. S4), cell numbers increased in the strains from Kiel, whereas in the Bornholm strains this was mainly observed on carbohydrates and carboxylic acids (Fig **5a**, Table S9 (TI), Effect of “Region*Carbon Group” F_5_ =3.798, p=0.002). At mid and late exponential phase, there were fewer sources on which cell numbers increased in the Kiel strains and there was no further increase in cell numbers on carbon sources in the Bornholm strains (Fig **5b**, Table S9 (TI), Effect of “Timing” F_2_ =67.528, p<0.0001). In the Kiel strains, there was still use of amines and carbohydrates at mid exponential and phenolic compounds in the late exponential phase (Fig **5a**, Table S9 (TI), Effect of “Timing*Region” F_2_ =9.290, p=0.0001). In addition, at 15°C compared to 22°C there were more carbon groups used in both Kiel and Bornholm strains (Fig **5b**, Table S9 (TE), Effect of “Temperature” F_1_ =37.202, p<0.0001), however differently so between the regions (Fig. **5b**, Table S9 (TE), Effect of “Temperature*Region” F_1_ =9.334, p=0.0023). In the Kiel strains resources from all groups except for the polymers lead to an increase in cell numbers and in Bornholm strains this was only the case on amines and carboxylic acids (Fig **5b**, Table S9 (TE), Effect of “Region*Carbon Group” F_5_ =3.798, p=0.002).

**Figure 5.**
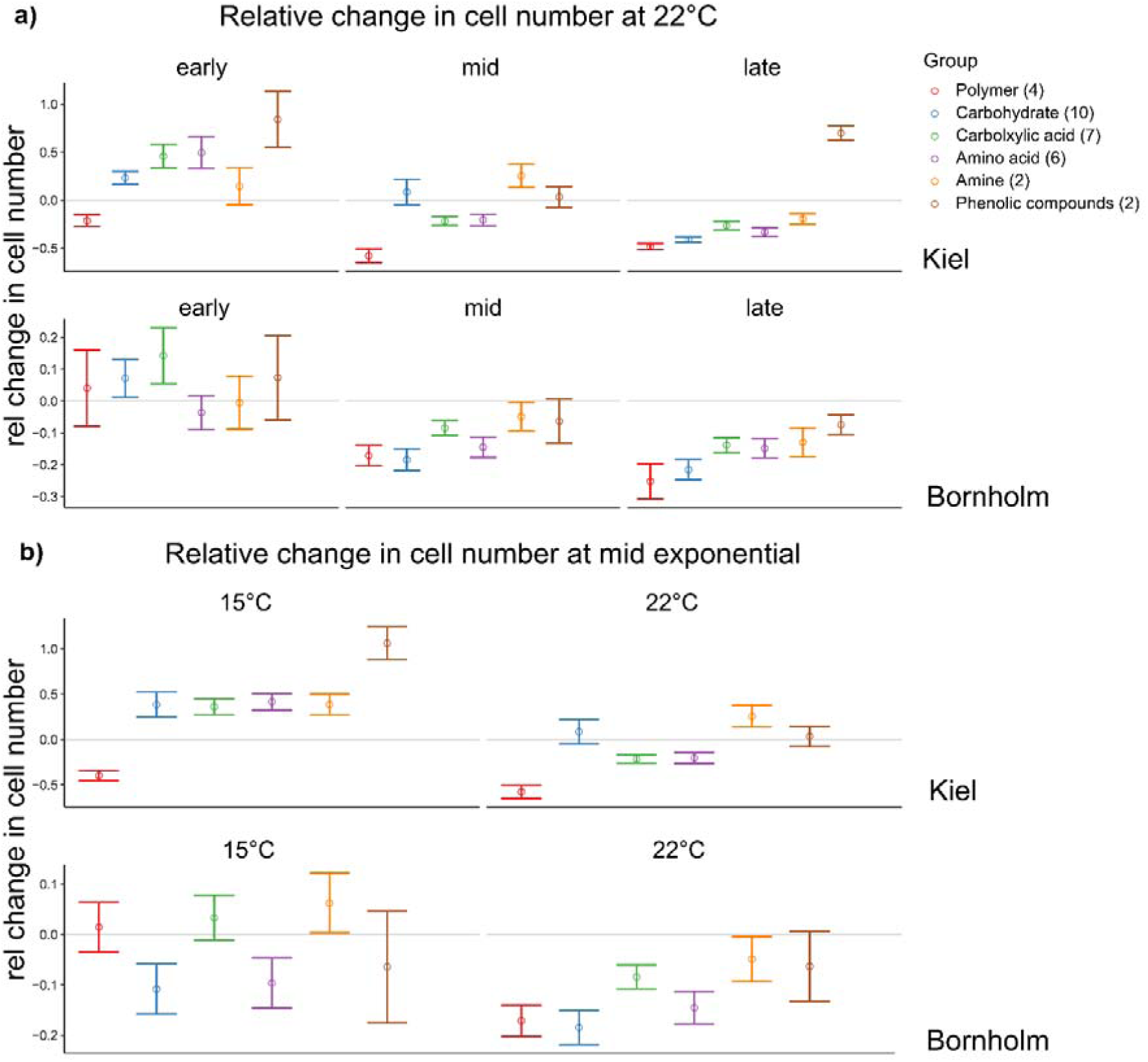
Relative change in cell numbers on the different carbon compound groups. The colours code for the different groups of carbon sources each source belongs to and on which we did the statistical analysis. Shown here are mean +/-1 SE for each group. However, each group contains different numbers of carbon sources (number in brackets) and a total of triplicate/source measurements of 4 or 2 strains for Kiel and Bornholm samples, respectively.

## Discussion

In our study on carbon uptake in *Ostreococcus sp*. from the Baltic Sea, we showed that there is an adaptive signature of warming or at the very least in the absence of molecular evidence, of processes acting on time-scales beyond acclimation and plasticity. Furthermore, we found different circumstances with respect to temperature and time in microbial growth curve when inorganic and organic carbon were used for growth.

With respect to the question if *Ostreococcus* has an adaptive signature of its origin we showed that strains from the warmer and more variable Kiel area showed higher growth rates in general and a more variable response to temperature as well. This was the opposite in the strains from the colder, less variable Bornholm Basin. These findings are consistent with the expectations based on short-term (within several generations) response measurements (Zhong *et al*., 2020) where the origin of the communities and thus their evolutionary history affected all key functional traits measured. Taking into account the wealth of theoretical work (Draghi & Whitlock, 2012; Botero *et al*., 2015; Ashander *et al*., 2016; Buckley & Kingsolver, 2019; Haaland & Botero, 2019) and experimental studies (Ketola & Saarinen, 2015; Schaum *et al*., 2016, 2018; Kristensen *et al*., 2018; Saarinen *et al*., 2018), we expected that the samples from the Kiel area will have been under selection in a more variable environment, giving rise to more variable responses with respect to growth.

Considering *how Ostreococcus* can adjust its growth mechanistically, we conclude that inorganic and organic carbon were taken up in different quantities depending on the stage of the microbial growth cycle and temperature to sustain positive growth that we observed throughout. At early exponential phase, both in strains from Kiel and Bornholm net primary production was below zero, which means that the carbon necessary to increase biomass could not have solely come from uptake of CO_2_ via photosynthesis. Rather, we found that cell numbers could increase on several organic carbon sources. This means, that likely at this early stage in the microbial growth curve, *Ostreococcus* sp. used organic carbon sources in the media to increase cell numbers. The organic carbon in the media could stem for example from the release of DOC by death of other *Ostreococcus* cells or bacteria (Thornton, 2014; Carlson & Hansell, 2015). At mid exponential and late exponential phase in the 22°C treatment, net primary production increased to above zero and the cell increase on carbon sources decreased to almost zero, indicating that at this stage in the microbial growth curve, growth was sustained mainly by uptake of CO_2_. However, in the 15°C treatment net primary production was still below zero and similarly, cells in the 15°C at mid exponential phase also increased on organic carbon sources compared to 22°C treatment. In summary, we found that when net primary production was lowest, the potential to grow on organic carbon sources was the highest and *vice versa* (see Fig. S5). Consequently, there seems to be a shift between the use of organic *vs* inorganic carbon leading to an increase cell numbers (Fig. S5). The origin of the strains further affected the likelihood and strength of this shift. In the samples that originated from the southern more thermally variable Kiel region, we found a stronger shift compared to the colder less variable Bornholm region (Fig S5). Several studies that have already investigated organic carbon uptake, found that indeed mixotrophy in microalgae increased biomass yields (for example (Kang *et al*., 2004; Liu *et al*., 2009; Pang *et al*., 2019)), however, these studies focused on optimising biofuel generation. It is still unclear, under what “natural” conditions microalgae preferably grow mixo-trophically or photo-trophically and what organic compounds may be available under natural conditions (Stickney *et al*., 2000; Flynn *et al*., 2013; Mitra *et al*., 2016).

Previous studies have already pointed towards evidence of *Ostreococcus* being able to sustain growth on sorbitol in the dark for circadian clock research (O-Neill *et al*., 2011; van Ooijen & Millar, 2012), but our study provides striking evidence that organic carbon sources are taken up readily in the light. This requires that we rethink our understanding of photoautotrophs and go beyond CO_2_ uptake. The consequences of the ability to take up organic carbon may be two-fold: on the one hand, the carbon pool used by *Ostreococcus* may not solely be in the form of CO_2_ but also DOM (dissolved inorganic matter). If, in general, many species of phytoplankton would indeed use other forms of carbon other than CO_2_, there might be a consequence on the carbon draw-down from the inorganic pool (Basu & Mackey, 2018). In particular, less DIC would be used directly for biomass production, but rather carbon would be taken up indirectly via the microbial shunt. On the other hand, using organic carbon sources puts the organisms in direct competition with other mixotrophic phytoplankton as well as heterotrophic organisms (e.g. bacteria). This second consequence is likely of more importance considering species interactions in microbial communities and thus ecosystem dynamics due to changes in the microbial loop (Meyer, 1994; Fenchel, 2008).

Generally, carbon acquisition *via* photosynthesis is cheap (Raven, 1991; Raven & Johnston, 1991) which is why there could be other reasons wherefore organic carbon is readily taken up by *Ostreococcus* under certain conditions. For example, the uptake of organic carbon compounds could be a “cheap” acquisition of organic nutrients (e.g. nitrogen, phosphorus) that are otherwise expensive to produce or acquire. The reduction of nitrate to organically available nitrogen (the same goes for phosphorous) is energy consuming (Timmermans *et al*., 1994). And at times where photosynthetic activity is low (i.e. at early and late exponential phase or lower temperatures), the available energy for such chemical conversions is low as well. As a result, using organic carbon compounds may be a way for the organism to acquire organic nutrients in a cheap way and use them for biomass formation and growth. Even if the growth on organic carbon compounds is not a consequence of requiring more carbon but rather organic nutrients, the effect this can have on competition that we highlighted above, may be similar. Whether the sources we tested were an organic carbon or organic nutrient source, could be investigated *via* the addition of DOC (dissolved organic carbon) or DOP (dissolved organic phosphorous) or DON (dissolved organic nitrogen) in manipulative experiments. The uptake of dissolved organic nutrients could then in addition be traced via mass spectrometry or HPLC (see for example Yan *et al*., 2012).

In this study, the growth on the organic carbon sources was not measured directly in culture, but rather as a potential to use a given source (ecoplates) (see Methods for details). Therefore, a manipulative experiment proving that the addition of an organic carbon source directly to the experimental culture increases growth, would be the next logical step. In addition, testing the effect of additional organic carbon sources on other functional groups of phytoplankton and heterotrophic organisms is necessary to characterize the consequence of possible competition between phototrophic and heterotrophic microbial species and how ecological dynamics would be affected.

In conclusion, we found that a small pico-phytoplankton species from the Baltic Sea does have an adaptive response to environmental change due to differences in ecological variability and evolutionary history. However, it is important to understand how growth as a response is mechanistically increased, as the differences in the carbon uptake related strategies may have implications on ecosystem dynamics and how well an organism can persist in future environments.

## Acknowledgements

We would like to thank Margarethe Nowicki, Richard Klinger, Jens-Peter Hermann, and the captain and crew of RV ALKOR for support at sea (cruises AL505 and AL507), and Stefanie Schnell for technical assistance in the laboratory at IMF Hamburg. This project was funded through a grant by Universitaet Hamburg to ES. Research was also partially funded by the Deutsche Forschungsgemeinschaft (DFG, German Research Foundation) under Germany‘s Excellence Strategy — EXC 2037 ‘CLICCS - Climate, Climatic Change, and Society’ — Project Number: 390683824, contribution to the Center for Earth System Research and Sustainability (CEN) of Universität Hamburg”.

## Author contributions

LL, FK and NM carried out the experiments. LL and ES retrieved, prepared, and maintained the phytoplankton cultures. LL and ES conceived and designed the experiment, supervised laboratory work and LL handled data analysis. LL wrote the first draft of the manuscript. All authors contributed equally to writing subsequent versions of the manuscript.

