## Supplementary Material for "Carbon acquisition in a Baltic pico-phytoplankton species - Where does the carbon for growth come from?"

Supplementary material to Carbon assimilation Listmann et al

Supplementary Figures

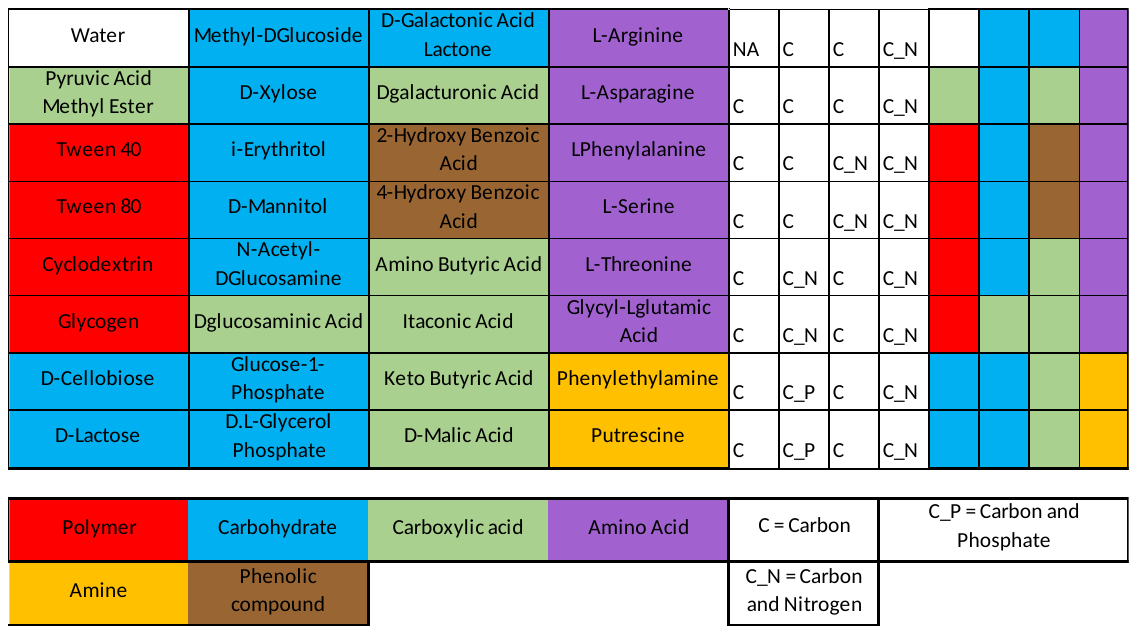

**Figure S1**: Sources for ecoplates (adjusted from Biolog EcoPlate^TM^ manual). Each source is present in triplicate such that there are four sources per row. The numbers of the sources are of use for Figure S4. The groups of organic sources that each source belongs to are shown below. We show the main nutrients in addition to carbon contained in each source.

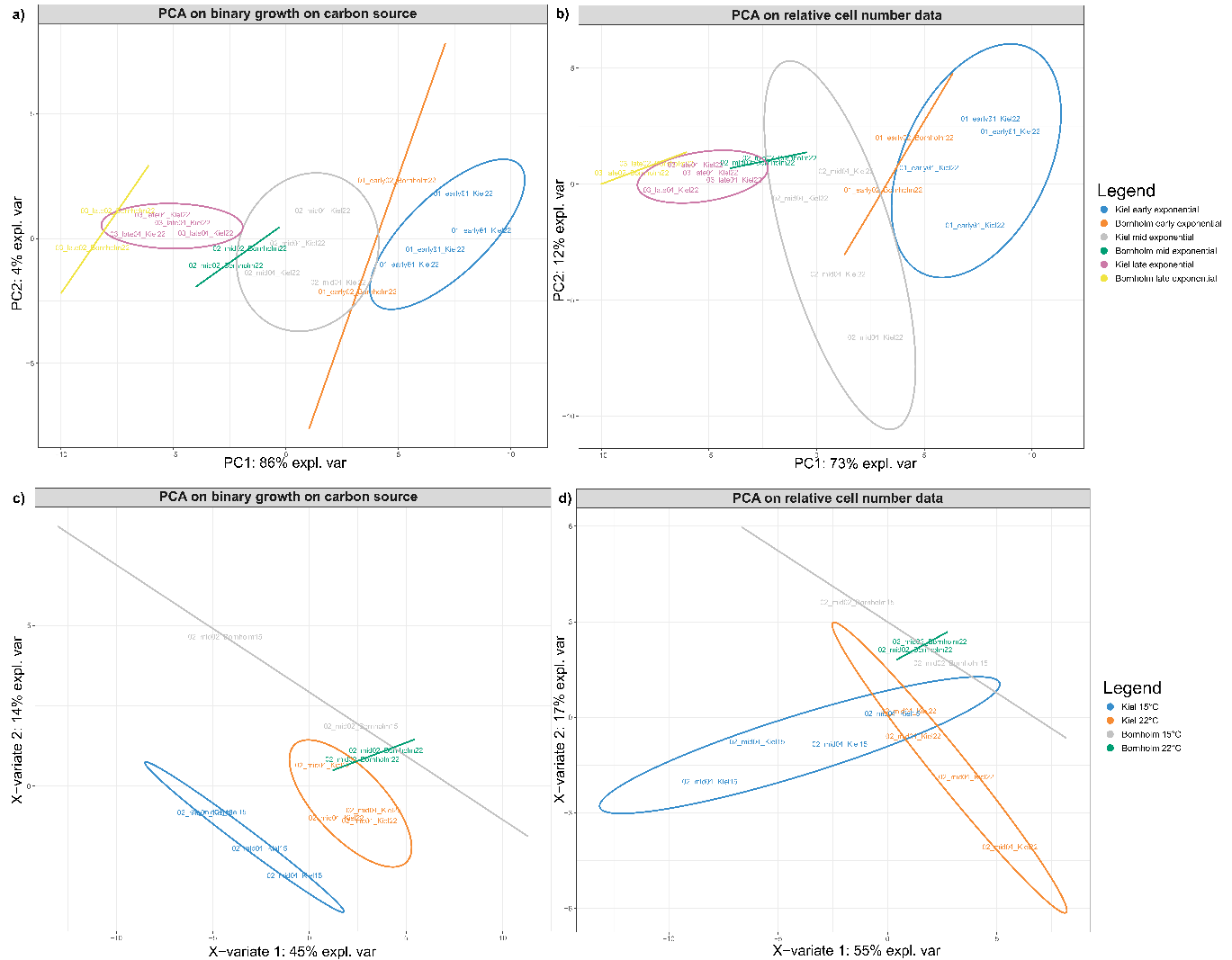

**Figure S2:** Panels **a)** and **b)** show the PCA results (based on grouping of regions and time points) for binary data (**a**) and relative data (**b**) of growth on the different carbon sources, respectively. Panels **c)** and **d)** show the PCA results (based on grouping of regions and temperature) at mid exponential phase for binary data (**c**) and relative data (**d**) of growth on the different carbon sources, respectively. The eclipses show the difference in organic carbon uptake between the regions and time points and temperatures at mid-exponential phase when the cultures were measured. A permanova (Table S4) underlines these results.

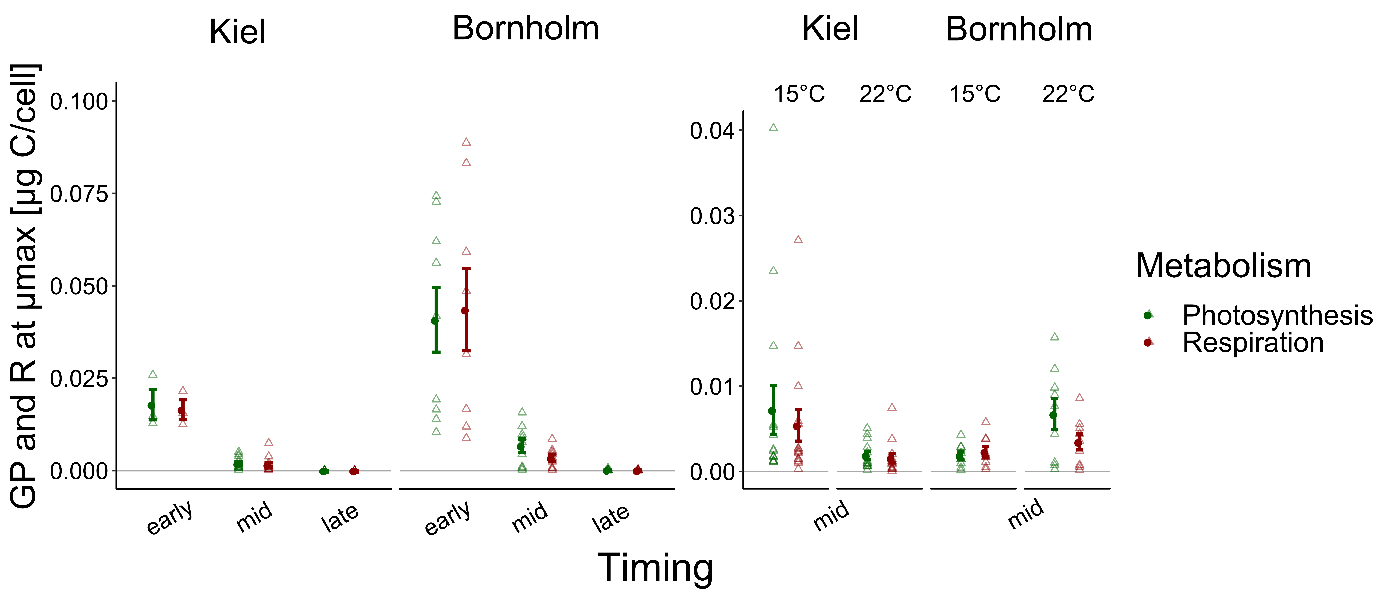

**Figure S3**: Photosynthesis and Respiration rates over Time at 22 °C (left panel) and at mid exponential phase in 15° and 22°C (right panel). Red shows the rates of gross photosynthesis whereas dark green shows the rate of respiration. Clear triangles show the measurements of each replicate or the strains. Circles and lines here are mean +/- 1 SE of n=15 and n=9 for Kiel and Bornholm samples, respectively

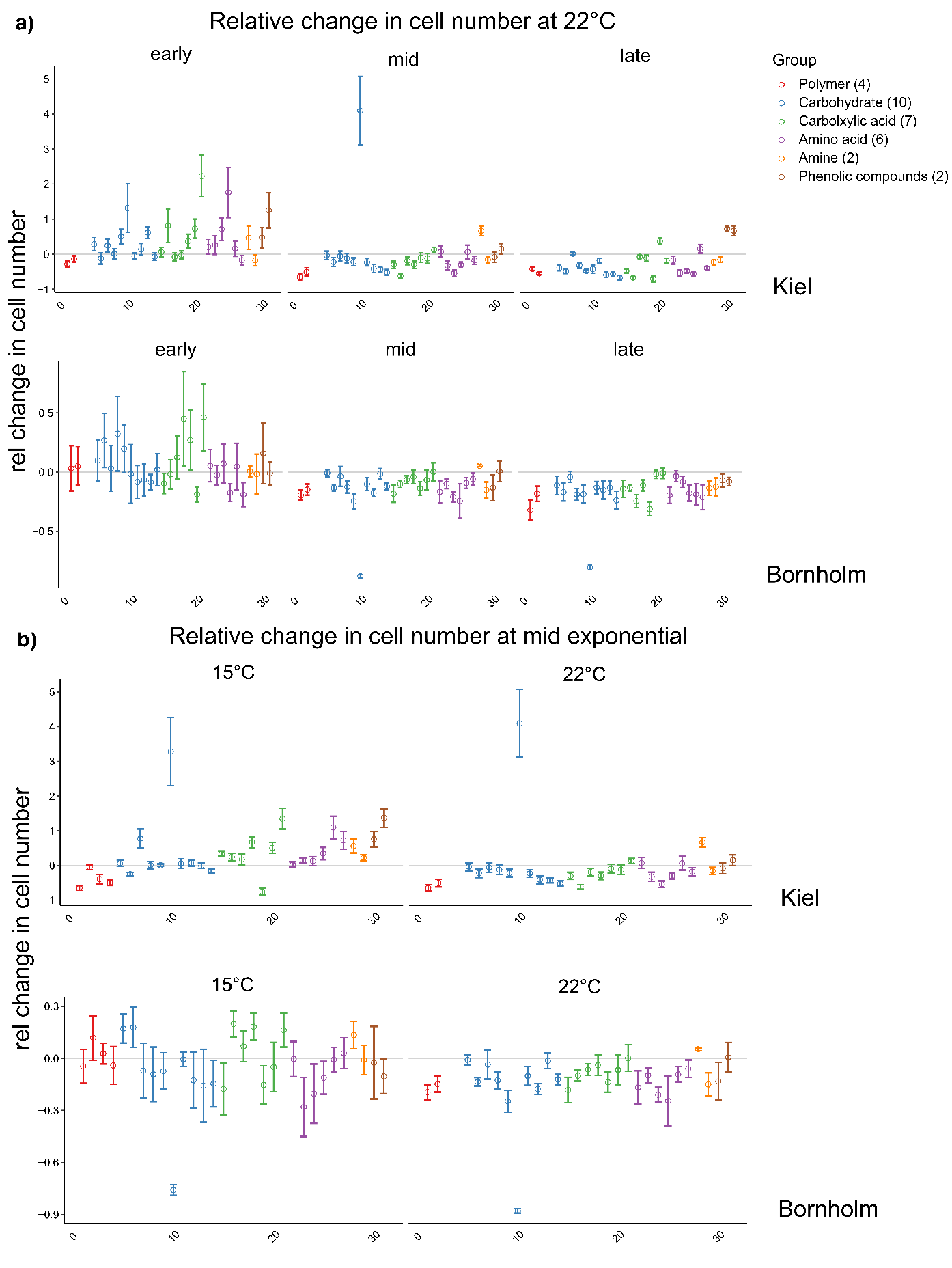
**Figure S4**: Relative change in cell numbers on the different carbon sources (for numbers see Fig. S1). The colours code for the different groups of carbon groups that each source belongs to. In panel **a**) the measurements at 22°C during the microbial growth curve are shown from early, mid and late exponential phase. In panel **b**) the measurements at mid exponential phase in 15°C and 22°C are shown.

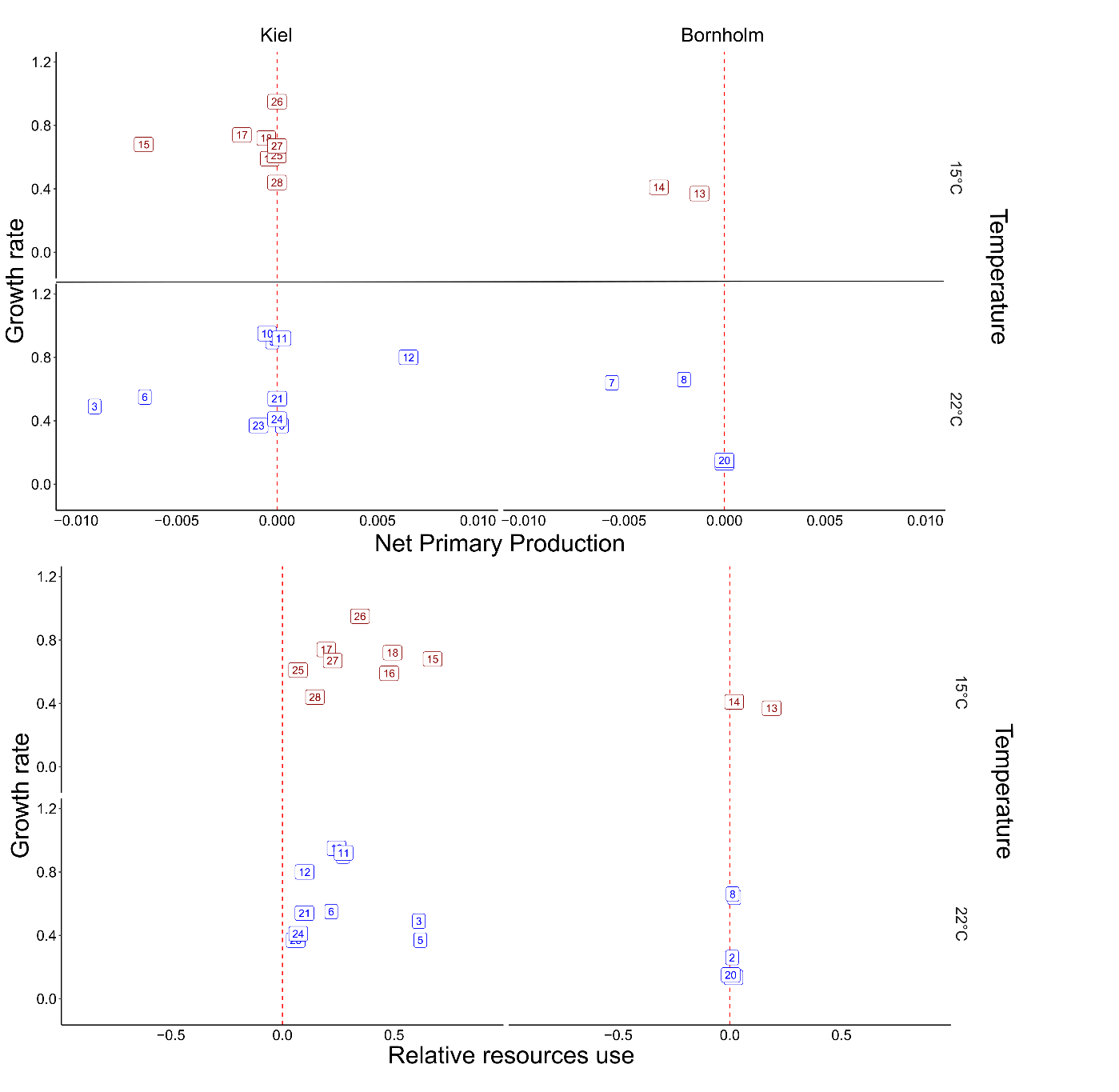

**Figure S5**: Growth rate calculated at early, mid and late exponential phase compared to the net primary production (top panel) and relative use of organic carbon (lower panel) for *Ostreococcus* grown at 15°C (blue) and 22°C (red). Theoretically, growth should not be possible below a net primary production of zero, meaning that in cultures that lie to the left side of the dashed red line in the top panel, another carbon source is necessary. In this study, we found that organic carbon is readily taken up seen in the lower panel for all cultures that lie to the right side of the dashed red line. Remarkably, there is a tendency of the samples with lowest net production to show a high relative resource use vice versa. See for example id. 6 and 3 in 15°C and id. 15 and 17 in 22°C both from the Kiel area.

Supplementary Tables

**Table S1:** Maximum growth rates and day at µmax calculated via curve fitting (see methods for details).

| Strain Name | Assay Temperature | BioRep | Experiment | µmax | Day at µmax | Region |
| --- | --- | --- | --- | --- | --- | --- |
| AL505_St19.1 | 15 | 2 | Exp3 | 0.501 | 9.483 | Bornholm |
| AL505_St19.1 | 15 | 3 | Exp3 | 0.485 | 9.772 | Bornholm |
| AL505_St19.1 | 22 | 1 | Exp3 | 0.660 | 8.931 | Bornholm |
| AL505_St19.1 | 22 | 2 | Exp3 | 0.561 | 8.818 | Bornholm |
| AL505_St19.1 | 22 | 3 | Exp3 | 0.543 | 8.992 | Bornholm |
| AL505_St19.5 | 15 | 1 | Exp3 | 0.498 | 9.530 | Bornholm |
| AL505_St19.5 | 15 | 2 | Exp3 | 0.512 | 9.288 | Bornholm |
| AL505_St19.5 | 15 | 3 | Exp3 | 0.546 | 8.928 | Bornholm |
| AL505_St19.5 | 22 | 1 | Exp3 | 0.548 | 8.968 | Bornholm |
| AL505_St19.5 | 22 | 2 | Exp3 | 0.540 | 8.832 | Bornholm |
| AL505_St19.5 | 22 | 3 | Exp3 | 0.586 | 8.293 | Bornholm |
| AL505_St19.9 | 15 | 1 | Exp2 | 0.454 | 11.832 | Bornholm |
| AL505_St19.9 | 15 | 2 | Exp2 | 0.440 | 12.149 | Bornholm |
| AL505_St19.9 | 22 | 1 | Exp2 | 0.475 | 11.208 | Bornholm |
| AL505_St19.9 | 22 | 2 | Exp2 | 0.512 | 10.487 | Bornholm |
| AL505_St21.1 | 15 | 1 | Exp1 | 0.524 | 9.182 | Kiel |
| AL505_St21.1 | 15 | 2 | Exp1 | 0.522 | 9.129 | Kiel |
| AL505_St21.1 | 22 | 1 | Exp1 | 0.762 | 7.732 | Kiel |
| AL505_St21.1 | 22 | 2 | Exp1 | 0.727 | 7.989 | Kiel |
| AL505_St21.1 | 22 | 3 | Exp1 | 0.808 | 7.503 | Kiel |
| AL505_St21.2 | 15 | 2 | Exp3 | 0.650 | 7.703 | Kiel |
| AL505_St21.2 | 15 | 3 | Exp3 | 0.648 | 7.746 | Kiel |
| AL505_St21.2 | 22 | 1 | Exp3 | 0.862 | 7.045 | Kiel |
| AL505_St21.2 | 22 | 2 | Exp3 | 0.729 | 7.563 | Kiel |
| AL505_St21.2 | 22 | 3 | Exp3 | 0.687 | 7.655 | Kiel |
| AL505_St21.3 | 15 | 1 | Exp3 | 0.722 | 7.236 | Kiel |
| AL505_St21.3 | 15 | 2 | Exp3 | 0.719 | 7.264 | Kiel |
| AL505_St21.3 | 22 | 1 | Exp3 | 0.738 | 7.237 | Kiel |
| AL505_St21.3 | 22 | 2 | Exp3 | 0.784 | 6.883 | Kiel |
| AL505_St21.4 | 15 | 1 | Exp2 | 0.587 | 8.434 | Kiel |
| AL505_St21.4 | 15 | 2 | Exp2 | 0.598 | 8.412 | Kiel |
| AL505_St21.4 | 15 | 3 | Exp2 | 0.615 | 7.978 | Kiel |
| AL505_St21.4 | 22 | 1 | Exp2 | 0.641 | 8.075 | Kiel |
| AL505_St21.4 | 22 | 2 | Exp2 | 0.644 | 7.966 | Kiel |
| AL505_St04.3 | 15 | 2 | Exp1 | 0.529 | 10.242 | Kiel |
| AL505_St04.3 | 15 | 3 | Exp1 | 0.545 | 9.844 | Kiel |
| AL505_St04.3 | 22 | 1 | Exp1 | 0.555 | 9.648 | Kiel |
| AL505_St04.3 | 22 | 2 | Exp1 | 0.551 | 9.773 | Kiel |
| AL505_St04.3 | 22 | 3 | Exp1 | 0.560 | 9.655 | Kiel |

**Table S2**: For the growth analysis in Biolog EcoPlate^TM^ the different strains were inoculated at different times of the batch cycle based on calculated day at µmax (see Table S1).

| Strain Name | Assay Temperature | Day of inoculation Early Exp | Day of inoculation Mid Exp | Day of inoculation Late Exp |
| --- | --- | --- | --- | --- |
| AL505_St21.1 | 22 | 3 | 7 | 11 |
| AL505_St21.2 | 22 | 3 | 7 | 11 |
| AL505_St21.3 | 22 | 3 | 7 | 11 |
| AL505_St21.4 | 22 | 3 | 7 | 11 |
| AL505_St04.3 | 22 | 3 | 7 | 11 |
| AL505_St19.1 | 22 | 7 | 11 | 15 |
| AL505_St19.5 | 22 | 7 | 11 | 15 |
| AL505_St21.1 | 15 | NA | 10 | NA |
| AL505_St21.2 | 15 | NA | 10 | NA |
| AL505_St21.3 | 15 | NA | 10 | NA |
| AL505_St21.4 | 15 | NA | 10 | NA |
| AL505_St04.3 | 15 | NA | 10 | NA |
| AL505_St19.1 | 15 | NA | 14 | NA |
| AL505_St19.5 | 15 | NA | 14 | NA |

**Table S3:** Here the results are shown of a permanova analysis using a Euclidean distance matrix on relative growth data on Biolog EcoPlate^TM^.

| Effect of Timing (TI) | | | | | | Effect of Temperature (TE) | | | | | |
| --- | --- | --- | --- | --- | --- | --- | --- | --- | --- | --- | --- |
| **Permanova** | **df** | **MS** | **F Model** | **R^2^** | **P-Value** | **Permanova** | **df** | **MS** | **F-value** | **R^2^** | **P-Value** |
| Timing | 2 | 243.292 | 19.558 | 0.681 | 0.001 | Temperature | 1 | 50.299 | 3.280 | 0.153 | 0.055 |
| Region | 1 | 54.885 | 4.412 | 0.077 | 0.026 | Region | 1 | 146.540 | 9.555 | 0.446 | 0.001 |
| Timing x Region | 2 | 11.79 | 0.948 | 0.033 | 0.431 | Temperature x Region | 1 | 9.342 | 0.609 | 0.028 | 0.575 |
| Residual | 12 |  |  |  |  | *Residual* | 8 |  |  |  |  |

**Table S4:** Here the results are shown of a PERMANOVA analysis using an Euclidean distance matrix on binary data with positive or no growth.

| Effect of Timing (TI) | | | | | | Effect of Temperature (TE) | | | | | |
| --- | --- | --- | --- | --- | --- | --- | --- | --- | --- | --- | --- |
| **Permanova** | **df** | **MS** | **F Model** | **R^2^** | **P-Value** | ***Permanova*** | **df** | **MS** | **F-value** | **R^2^** | **P-Value** |
| Timing | 2 | 228.278 | 35.576 | 0.781 | 0.001 | Temperature | 1 | 52.500 | 7.706 | 0.244 | 0.008 |
| Region | 1 | 42.389 | 6.606 | 0.073 | 0.015 | Region | 1 | 102.333 | 15.021 | 0.476 | 0.002 |
| Timing x Region | 2 | 4.139 | 0.645 | 0.014 | 0.573 | Temperature x Region | 1 | 5.500 | 0.807 | 0.026 | 0.419 |
| Residual | 12 |  |  |  |  | Residual | 8 |  |  |  |  |

**Table S5:** Analysis report for growth rate: linear mixed effects model was used for the effect Timing and Region and its interaction as well as the effect of Temperature and Region at mid exponential phase were performed. The best model was selected via AICc analysis with a minimal difference of 2. We analysed first the whole data set via a global model and then the two regions separately to also investigate the differences in growth between the strains.

| Global model | | | |  | | | |
| --- | --- | --- | --- | --- | --- | --- | --- |
| **Effect of Timing (TI)** | | | | **Effect of Temperature (TE)** | | | |
| **Anova of lme** | **df** | **F-value** | **P-Value** | **Anova of lme** | **df** | **F-value** | **P-Value** |
| Timing | 2 | 4.990 | 0.011 | Temperature | 1 | 15.430 | 0.0005 |
| Region | 1 | 11.779 | 0.001 | Region | 1 | 40.162 | <0.0001 |
|  |  |  |  | Temperature x Region | 1 | 0.353 | 0.557 |

| Regional model Kiel | | | | *Regional model Bornholm* | | | |
| --- | --- | --- | --- | --- | --- | --- | --- |
| **Anova of lme** | **df** | **F-value** | **P-Value** | **Anova of lme** | **df** | **F-value** | **P-Value** |
| Temperature | 1 | 31.725 | 0.0001 | Temperature | 1 | 7.560 | 0.029 |
| Isolate | 4 | 7.618 | 0.003 | Isolate | 1 | 0.047 | 0.833 |
| Temperature x Isoalte | 4 | 6.426 | 0.005 | Temperature x Isolate | 1 | 1.372 | 0.280 |

**Table S6:** Analysis report for net primary production: linear mixed effects model was used for the effect Timing and Region and its interaction as well as the effect of Temperature and Region at mid exponential phase were performed. The best model was selected via AICc analysis with a minimal difference of 2.

| Global model | | | |  | | | |
| --- | --- | --- | --- | --- | --- | --- | --- |
| **Effect of Timing (TT)** | | | | **Effect of Temperature (TE)** | | | |
| **Anova of lme** | **df** | **F-value** | **P-Value** | **Anova of lme** | **df** | **F-value** | **P-Value** |
| Timing | 2 | 36.583 | <.0001 | Temperature | 1 | 8.818 | 0.0005 |
| Region | 1 | 0.620 | 0.431 | Region | 1 | 1.299 | 0.261 |
| Timing x Region | 2 | 5.093 | 0.006 | Temperature x Region | 1 | 0.019 | 0.891 |

**Table S7:** Analysis report for growth on organic carbon: linear mixed effects model was used for the effect Timing and Region and its interaction as well as the effect of Temperature and Region at mid exponential phase were performed. The best model was selected via AICc analysis with a minimal difference of 2.

| Global model | | | |  | | | |
| --- | --- | --- | --- | --- | --- | --- | --- |
| **Effect of Timing (TI)** | | | | **Effect of Temperature (TE)** | | | |
| **Anova of lme** | **df** | **F-value** | **P-Value** | **Anova of lme** | **df** | **F-value** | **P-Value** |
| Timing | 2 | 12.398 | 0.002 | Temperature | 1 | 5.730 | 0.062 |
| Region | 1 | 6.531 | 0.063 | Region | 1 | 11.416 | 0.027 |

**Table S8:** Analysis report for correlation analysis between growth on organic carbon and net primary production. The analysis was done separately for the data from the two regions.

| Correlation Kiel | | | | *Correlation Bornholm* | | | |
| --- | --- | --- | --- | --- | --- | --- | --- |
| **Cor.test** | **df** | **t-value** | **P-Value** | **Cor.test** | **df** | **t-value** | **P-Value** |
|  | 17 | -2.445 | 0.026 |  | 6 | -3.435 | 0.0139 |

**Table S9:** Analysis report for relative change in cell numbers: linear mixed effects model was used for the effect Timing, Region and Carbon Group and its interaction as well as the effect of Temperature, Region and Carbon Group at mid exponential phase. The best model was selected via AICc analysis with a minimal difference of 2.

| Global model | | | |  | | | |
| --- | --- | --- | --- | --- | --- | --- | --- |
| **Effect of Timing (TT)** | | | | **Effect of Temperature (TE)** | | | |
| **Anova of lme** | **df** | **F-value** | **P-Value** | **Anova of lme** | **df** | **F-value** | **P-Value** |
| Timing | 2 | 67.528 | <.0001 | Temperature | 1 | 37.202 | <.0001 |
| Region | 1 | 0.213 | 0.668 | Region | 1 | 2.921 | 0.162 |
| Carbon Group | 5 | 10.522 | <.0001 | Carbon Group | 5 | 8.608 | <.0001 |
| Timing x Region | 2 | 9.290 | 0.0001 | Temperature x Region | 1 | 9.334 | 0.0023 |
| Region x Carbon Group | 5 | 3.798 | 0.002 | Region x Carbon Group | 5 | 5.007 | 0.0002 |
